# DENOISING: Dynamic Enhancement and Noise Overcoming in Multimodal Neural Observations via High-density CMOS-based Biosensors

**DOI:** 10.1101/2024.05.29.596467

**Authors:** Xin Hu, Brett Addison Emery, Shahrukh Khanzada, Hayder Amin

## Abstract

Large-scale multimodal neural recordings on high-density biosensing microelectrode arrays (HD-MEAs) offer unprecedented insights into the dynamic interactions and connectivity across various brain networks. However, the fidelity of these recordings is frequently compromised by pervasive noise, which obscures meaningful neural information and complicates data analysis. To address this challenge, we introduce DENOISING, a versatile data-derived computational engine engineered to adjust thresholds adaptively based on large-scale extracellular signal characteristics and noise levels. This facilitates the separation of signal and noise components without reliance on specific data transformations. Uniquely capable of handling a diverse array of noise types (electrical, mechanical, and environmental) and multidimensional neural signals, including stationary and non-stationary oscillatory local field potential (LFP) and spiking activity, DENOISING presents an adaptable solution applicable across different recording modalities and brain networks. Applying DENOISING to large-scale neural recordings from mice hippocampal and olfactory bulb networks yielded enhanced signal-to-noise ratio (SNR) of LFP and spike firing patterns compared to those computed from raw data. Comparative analysis with existing state-of-the-art denoising methods, employing SNR and root mean square noise (RMS), underscores DENOISING’s performance in improving data quality and reliability. Through experimental and computational approaches, we validate that DENOISING improves signal clarity and data interpretation by effectively mitigating independent noise in spatiotemporally structured multimodal datasets, thus unlocking new dimensions in understanding neural connectivity and functional dynamics.

## 1 Introduction

The intricate exploration of computational neural dynamics investigated through the coordinated activity of interconnected neural populations, especially within the hippocampus and olfactory bulb, has been a cornerstone of contemporary neuroscience research (1). These regions, central to spatial contextual learning, episodic memory, and olfactory processing, demonstrate remarkable neuroplasticity and are key to understanding the complex interplay of neural circuits in cognitive functions (2,3). The hippocampus, a hub for information flow and synaptic plasticity, is crucial for the formation and retrieval of memories. Its ability to undergo structural and functional modifications in response to stimuli underscores the dynamic nature of neural networks (4). Similarly, the olfactory bulb (OB) serves as the initial stage of olfactory processing, transforming odorant signals into neural representations through its intricate layers and diverse neuronal interactions. The regions’ unique capacity for adult neurogenesis offers a window into the mechanisms underlying sensory perception and memory integration (5,6).

Extracellular neural recordings have long been a fundamental tool in neuroscience, offering insights into the electrical activity of neurons in their native environment (7). The evolution of microelectrode arrays (MEAs) has expanded the scope of these observations, facilitating the simultaneous recording of multiple neural signals. Among the MEA technologies, high-density CMOS-based biosensing platforms (HD-MEAs) stand out due to their unparalleled technical capabilities (8,9). With the capacity to feature thousands of electrodes, these arrays can simultaneously capture a detailed panorama of neural activity across extensive networks, providing a dense sampling of electrical signals with high spatial and temporal resolution. This dense array structure allows for an in-depth analysis of neural interactions, offering a window into the synchronous and asynchronous patterns that underlie dynamical processes and functional connectome in multimodal neural networks and circuits (10–14). This technology has facilitated a shift from the study of isolated neural pathways to an integrative view of brain’s functional networks, bridging gaps in our knowledge of how neuronal ensembles coordinate to produce complex behaviors and cognitive functions (15). HD-MEAs significantly enhance brain slice studies, merging *ex vivo* biosensing precision with brain tissue complexity. This approach allows for detailed exploration of electrical spiking activity and rhythmic dynamics of local field potentials (LFPs) under controlled conditions, thus enabling researchers to investigate environmental factors, apply pharmacological agents, or introduce genetic modifications to elucidate their effects on neural activity (10,11,14,16–18).

Despite advancements in these electrophysiological technologies, capturing the full spectrum of neural patterns within these complex networks remains challenging. The fidelity of extracellular neural recordings is frequently susceptible to a range of independent noise sources, including electrical interference from the recording equipment, mechanical vibrations caused by external or internal laboratory factors, and environmental noise, such as electromagnetic fields. Removing noise from extracellular neural recordings poses several challenges due to the complex nature of neural signals and the non-stationary characteristics of noise. Neural signals often exhibit irregular firing patterns and non-Gaussian distributions, while noise can vary in amplitude and frequency content over time (19). Additionally, the presence of overlapping signals from multiple neuronal ensembles further complicates the task of noise removal. Several classical denoising strategies typically focused on temporal, spatial, or transform domains (i.e., Wavelet or Fourier) (20), which often fall short due to oversimplified assumptions about signal and noise characteristics (21,22). The limited adaptability of these methods inadequately addresses the complexity and heterogeneity in large-scale neural recordings. This limitation not only impedes practical data analysis but also restricts our understanding of essential neural mechanisms.

Moreover, unsupervised denoising methods may introduce additional bias, as they might inadvertently emphasize or suppress certain signal features without ground truth verification, potentially leading to incorrect scientific inferences by distorting the underlying neural processes (23). Despite existing classical methods for denoising, current strategies do not effectively address the unique challenges posed by large-scale neural recordings captured by HD-MEAs. This gap underscores the critical need for new denoising approaches designed explicitly for HD-MEA data, which can dynamically adapt to its complexity, minimizing bias and significantly enhancing signal clarity for robust and accurate neural analysis.

In response to these challenges, we introduce DENOISING, a computational framework developed to transcend the limitations of conventional denoising techniques. Leveraging insights from recent studies highlighting the intricate dynamics and plasticity within the hippocampus and OB, DENOISING employs an adaptive engine to enhance the clarity and reliability of multidimensional neural recordings. Our method dynamically adjusts to the specific spatiotemporal characteristics (i.e., firing pattern statistics, network synchrony, burst, and waveform shapes) of extracellular signals and independent noise, facilitating a more nuanced separation of signal and noise components. This approach is predicated on a deep understanding of the spatial and temporal structures of neural activity, informed by the complex interplay of vast neuronal ensembles within these critical brain regions.

In the following sections, we detail our method’s effectiveness through its use in analyzing large-scale neural LFP and spike data from complex hippocampal-cortical and olfactory networks. This includes demonstrating significant signal-to-noise ratio (SNR) (24) enhancement, mapping topographical propagation features, classifying patterns based on their initiation and transmission, and waveform characteristics. Furthermore, we benchmark DENOISING against traditional methods, demonstrating our approach’s capacity to enhance data quality and reliability.

Our study illuminates the path toward more accurate and comprehensive analyses of HD-MEA’s extracellular recordings, highlighting the potential to unlock new dimensions in our understanding of large-scale neural connectivity and functional dynamics and opening new avenues for exploring the mechanisms of learning, memory, and sensory processing.

## 2 MATERIALS AND METHODS

### 2.1 DENOISING Framework

We employed an adaptive waveform-based thresholding technique designed specifically for processing signals captured by high-density CMOS-based microelectrode arrays. This involves setting customized thresholds for noise removal based on multiple signal waveform characteristics, including amplitude variations, frequency content, and waveform shape irregularities. The thresholding process is dynamically adjusted in real-time, leveraging the dense data acquisition capabilities of the arrays to detect and isolate noise components from true neural signals. The operational framework employed a composite of these signal features to define noise versus signal criteria, which are applied instantaneously to each waveform detected across the array’s multiple recording channels. This method ensures a robust noise reduction while preserving the integrity of the biological signal, making it highly suitable for environments with variable noise conditions often encountered in high-density array recordings.

### 2.2 Animals and Acute Brain Slice Preparation

Our study utilized 12-week-old female C57BL/6j mice (Charles River Laboratories, Germany), and ensured all procedures complied with European and national animal welfare regulations (Tierschutzgesetz), with approval license (Landesdirektion Sachsen; 25–5131/476/14). Brain slices were prepared according to our previous report (10,11). Following anesthesia with 0.05% inhaled isoflurane (Primal, Germany), mice were decapitated, and their brains were extracted and submerged in a chilled sucrose solution for slicing. Using a Leica Vibratome VT1200S (Leica Microsystems, Germany), we prepared 300 μm thick horizontal brain slices, cut at 0–2 °C in aCSF solution saturated with 95% O_2_ and 5% CO_2_ (pH = 7.2–7.4) of a high sucrose solution containing in mM: 250 Sucrose, 10 Glucose, 1.25 NaH_2_PO_4_, 24 NaHCO_3_, 2.5 KCl, 0.5 Ascorbic acid, 4 MgCl_2_, 1.2 MgSO_4_, 0.5 CaCl_2_. Furthermore, hippocampal and OB slices were incubated for 45 min at 32 °C and then allowed to recover for at least 1 hour at room temperature before recording with HD-MEAs in a recording aCSF solution containing in mM: 127 NaCl, 2.5 KCl, 1.25 NaH_2_PO_4_, 24 NaHCO_3_, 25 Glucose, 1.2 MgSO_4_, 2.5 CaCl_2_, and the solution was aerated with 95% O_2_ and 5% CO_2_.

### 2.3 Multimodal Extracellular Recordings and LFP/Spike Events Detection

Extracellular neural activity was recorded using HD-MEAs crafted from complementary-metal-oxide-semiconductor (CMOS) technology, coupled with a bespoke acquisition system (3Brain AG, Switzerland). The CMOS chip featured 4096 electrodes organized in a 64×64 array with a pitch of 42 μm, creating an active sensing area of approximately 7 mm^2^, an ideal dimension for comprehensive recordings from both hippocampal-entorhinal cortex and olfactory bulb (OB) tissues at 14 kHz/electrode sampling frequency (10,11). The on-chip amplification circuit allowed band-pass filtering from 1 Hz to 5 kHz, sufficient to record slow and fast neural activity. The hippocampal-entorhinal cortical recordings spanned six layers: dentate gyrus (DG), Hilus, CA3, CA1, entorhinal cortex (EC), and perirhinal cortex (PC). Similarly, OB recordings encompassed neuronal signals across five distinct layers: the olfactory nerve layer (ONL), glomerular layer (GL), external plexiform layer (EPL, we referred to as the projection layer), the olfactory cortex (OCx), and granule cell layer (GCL). Integration of a modular stereomicroscope (Leica Microsystems, Germany) allowed for simultaneous acute slice imaging and extracellular recording, facilitating the correlation of spatial tissue organization with electrode activity. Event detection for LFPs and multi-unit spiking activity (MUA) was conducted using commercial software (3Brain AG), where data was first refined by applying a low-pass filter (1– 100 Hz) for LFPs and a band-pass filter (300–3500 Hz) for MUA. Following the filtering, events were detected using hard thresholding alongside precise timing spike detection (PTSD) algorithms, respectively (10,11). This sequence ensured that the frequency components suitable for describing LFPs and spikes were accurately isolated before event detection, improving the specificity and accuracy of the detected events.

### 2.4 Topographical Spatiotemporal Voltage Maps, CATs, and Event Incidence

To assess the impact of DENOISING in enhancing dynamical spatiotemporal information in the hippocampus and OB subregional networks, we computed averaged LFP and spike event frequencies across their interconnected layers. By employing high-resolution, multimodal recordings, we generated dynamic topographical maps for LFP and spike data within respective 50 ms and 10 ms time bins. Illustrated in pseudo-color, these maps demonstrate the spatial distribution of electrical activity per event, accentuating the enhanced clarity of neuronal interactions after noise removal. Spatiotemporal activity propagation was quantified by analyzing the center of activity trajectories (CATs)(10). Furthermore, we determined long-range event incidence rates and their distributions, leveraging simultaneous recordings from extensive subnetworks to clarify decontaminated initiation sites and their propagation across layers after employing DENOISING.

### 2.5 Traditional Noise-removal Methods

DENOISING was evaluated against established noise reduction techniques in neural recordings, categorized into signal and transform domain methods. In the signal domain, we applied one-dimension causal forward-in-time FIR filtering (25) (https://docs.scipy.org/doc/scipy/reference/generated/scipy.signal.lfilter.html#scipy.signal.lfilter) and the Savitzky-Golay algorithm (26) for smoothing the signal by fitting a polynomial to a segment of neural data points via least squares regression (https://docs.scipy.org/doc/scipy/reference/generated/scipy.signal.savgol_filter.html). For transform domain denoising, we utilized Wavelet (WT) and Fourier Transform (FT) methods, which transform the recorded signals into different frequency components, where the noise removal or filtering operation is applied more effectively. WT denoising involves convolution with a wavelet function to isolate signal frequencies computed using the PyWavelets package (https://github.com/PyWavelets/pywt). FT approaches modify signal frequencies in the frequency domain before an inverse transformation (https://numpy.org/doc/stable/reference/generated/numpy.fft.ifft.html). These methodologies provide a foundation to demonstrate DENOISING’s performance extracting clean neural signals.

### 2.6 Extracellular Waveform Characterization and Clustering Assessment

We performed unsupervised clustering analysis to group similar waveform shapes of LFP patterns using principal component analysis (PCA) clustering with the mean-shift algorithm (27,28). PCA was applied to reduce the dimensionality of the waveform data while preserving the essential features (https://scikit-learn.org/stable/modules/generated/sklearn.decomposition.PCA.html). The mean-shift algorithm was then employed to identify clusters in the reduced-dimensional space (https://scikit-learn.org/stable/modules/generated/sklearn.cluster.MeanShift.html). Unlike other algorithms, such as k-means (29), mean-shift does not require the number of clusters to be specified, as it automatically determines the clusters based on data density. This approach allowed us to uncover distinct patterns of LFP activity within the hippocampal and OB slices in raw and denoised data.

To identify and cluster spiking activity from the recordings, we employed an unsupervised spike sorting algorithm compatible with large-scale neural recordings (30) and available on GitHub (https://github.com/mhhennig/HS2). The spike sorting algorithm was modified and implemented to extract various features, such as spike waveforms, spike amplitudes, and spike timing, to isolate individual spikes and group them into distinct clusters corresponding to different neuronal units estimated from multimodal large-scale neuronal ensembles. This process enabled us to differentiate between different types of firing electrodes and discern their spiking patterns within the hippocampal and OB slices.

Furthermore, we computed silhouette coefficients (SC) to assess the quality of the clustering results obtained from both LFP waveform shapes and multi-unit spiking activity(31). Silhouette coefficients measure the coherence and separation of clusters, providing a quantitative measure of clustering quality (https://scikit-learn.org/stable/auto_examples/cluster/plot_kmeans_silhouette_analysis.html). Higher silhouette coefficients indicate better-defined and more distinct clusters, while negative coefficients suggest overlapping or poorly separated clusters. By computing silhouette coefficients, we could objectively evaluate the effectiveness of our DENOISING method through clustering algorithms in capturing the underlying structure of the neural activity data compared to raw noise-contaminated data. This highlights the only actual partition of waveforms from the firing electrodes without the bias of the clustering algorithm used to obtain them.

### 2.7 Performance Benchmark Analysis

To evaluate the performance of DENOISING, we employed three metrics – a normalized signal-to-noise ratio difference (SNR_DR_) (24), SNR distribution (32)and root mean square noise (RMS) (33). SNR_DR_ offers a normalized measure of improvement, facilitating meaningful comparisons even when baseline SNR levels vary widely between recordings. This metric highlights the relative gains achieved by our denoising method, providing insights into its efficiency in enhancing signal quality relative to the initial noise level. SNR distribution evaluates the performance of the denoising method across the entire large-scale network with high channel counts. It demonstrates the method’s general applicability and effectiveness under diverse experimental conditions. RMS is a statistical measure that quantifies the magnitude of signal variation, assessing the effectiveness of noise reduction. This metric further quantifies improvements in signal quality, complementing the SNR metrics by offering a direct measure of the denoising impact on signal magnitude. The dynamic range of the SNR using the logarithmic decibel scale (SNR_S_) is defined as:

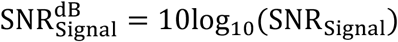

The normalized SNR_DR_ is defined as the normalized difference between the (SNR) of the denoised signal and the SNR of the raw signal, given as:

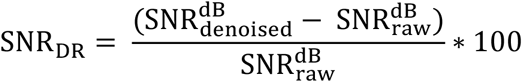

In addition, to determine the distribution of SNR in large-scale recordings, we computed SNR as the ratio of mean firing activity in all active channels to standard deviation across the time domain.

Furthermore, the RMS measures the magnitude of the varying components of biosignals. It provides a single number that represents the noise level in a way comparable across raw and denoised conditions. The RMS is defined as:

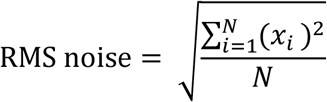

Where *x*_*i*_ denotes the individual values of the firing electrodes (signals), and *N* represents the number of data samples.

### 2.8 Data and Statistical Analysis

All analyses used in this study were developed and implemented with custom-written Python scripts available on our Lab’s GitHub (https://github.com/HayderAminLab/DENOISING). Any employed packages are cited accordingly. All statistical analyses were performed with Python and Originlab 2022. All data in this work were expressed as the mean ± standard error of the mean (SEM). Differences between groups were examined for statistical significance, where appropriate, using one-way analysis of variance (ANOVA) or Kolmogorov-Smirnov test followed by Tukey’s posthoc testing. *P < 0*.*05* was considered significant.

## 3 Results

### 3.1 DENOISING Principles

We implemented the DENOISING method as a novel approach for denoising extracellular large-scale recordings obtained from hippocampal and olfactory bulb slices. The primary objective of our method was to effectively remove noise while preserving the essential features of the signals, thereby enhancing the clarity and precision of LFP and spike patterns (**Figure 1**). The method is implemented through waveform-based thresholding, which operates directly on the time-domain representation of the signals. This method involves setting a threshold level based on the characteristics of the waveforms and removing signal segments below this threshold, which are considered noise. We separately applied waveform-based thresholding to both LFP (**Figures 1a-c**) and spike (**Figures 1d-f**) signals to ensure optimal denoising performance for each signal type. The DENOISING method utilized several identifiers derived from network-wide features to enhance the noise-removal process and improve its adaptability to different datasets. These identifiers were used to set up a template containing specific values of spatiotemporal pattern features, which could be cast off for testing with other datasets. The network features included - firing frequency (synchrony), number of firing electrodes (adapted to the structural clusters identified by optical imaging information combined with electrophysiology recordings), event duration (min-max range), event amplitude, and bursting frequency. Following the integration of denoising parameters and their automated application to the hippocampal and OB recorded datasets, the DENOISING method exhibited a substantial reduction in independent noise artifacts, as depicted in network-wide activity represented in 5-minute raster plots (**Figure 1**).

**Figure 1.**
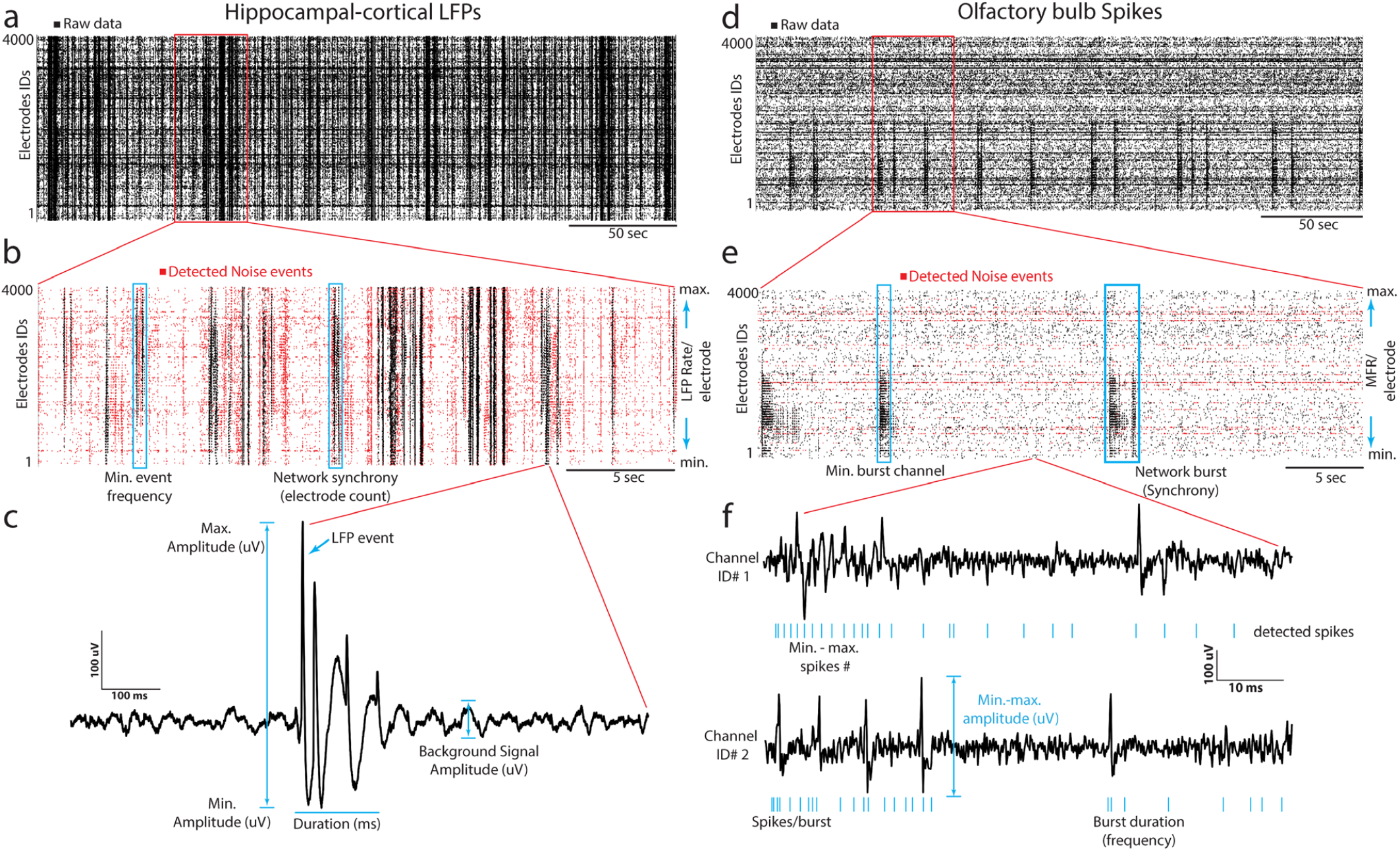
DENOISING Principles and Implementation. **a)** LFP signal rastergram from hippocampal-cortical recordings, illustrating waveform-based thresholding (black points indicate raw data with noise). **b)** Detailed rastergram highlighting noise reduction and preservation of essential signal features (red points indicate the detected noise channels after DENOISING). **c)** LFP waveform features are used in denoising to enhance clarity in spatiotemporal LFP events. **d-f)** Analogous to (a-c), for olfactory bulb spike recordings, showcasing spike-based feature application in denoising.

### 3.2 Validation with Multimodal Neural Recordings

To underscore the physiological validity of our neural recordings, we initially focused on exploiting multimodal neural recordings derived from extracellular LFP and spike signals within well-characterized hippocampal-cortical and olfactory bulb circuits (10,11). These systems are known for their intricate spatiotemporal dynamics, including specific initiation sites and subsequent propagation across distinct neural layers. Such detailed mapping is crucial for understanding the complex interplay of neural activities that underpin cognitive and sensory processing (1–3). We applied the DENOISING framework to process these recordings, intended not only to assess its noise reduction capabilities but also to demonstrate that our approach preserves the biological fidelity of these dynamic neural interactions.

Representative event traces from key regions of the hippocampal-cortical network regions (i.e., CA1, CA3, and EC) and OB (i.e., GL, PL, and GCL) were analyzed. These traces captured a diverse array of LFP and spike biosignal signatures both in their raw, unprocessed state and following noise reduction using DENOISING across multiple categories of noise examples (**Figure 2**). The significant denoising performance of our method for both LFP and multi-unit spiking-based patterns was demonstrated in - **i)** effectively discarded spatiotemporal segments of unwanted noise, even when the stringent hard threshold and PTSD algorithms detected events. This precision was evident in representative traces from CA1 and GL electrodes (**Figures 2a-b**), highlighting the method’s ability to discern genuine neural activity from noise. **ii)** identified and eliminated non-physiological signals contaminated with independent noise, including 50 Hz environmental noise (**Figure 2a**; EC electrode) and mechanical artifacts stemming from perfusion glitches (**Figure 2b**; PL electrode). This capability underscores the method’s robustness in preserving the fidelity of neural recordings. **iii)** suppressed inductive coupling noise originating from CMOS-chip electrical wiring before signal amplification. DENOISING enhanced the accuracy of neural signal interpretation by distinguishing and attenuating false signals with similar frequency and amplitude to neural oscillations, as evidenced by traces in (**Figures 2a-b**; CA3 and GCL electrodes). **iv)** removed falsely detected spurious spiking patterns embedded within the physiological spiking activity observed across electrodes in CA1, CA3, EC, and GL regions (**Figures 2a-b**). This ensures the extracted spiking activity reflects the true neuronal firing dynamics, facilitating more precise insights into neural circuit function and information processing within these critical brain regions. **v)** rectified false signal patterns that emerged from dysregulated chip calibration, as illustrated in traces from PL and GCL electrodes (**Figure 2b**). The auto-zeroing circuit integrated into the CMOS-chip, is regularly calibrated to the electrode’s DC voltage, which could be impaired due to various noise sources and light effects, leading to signal contamination.

**Figure 2.**
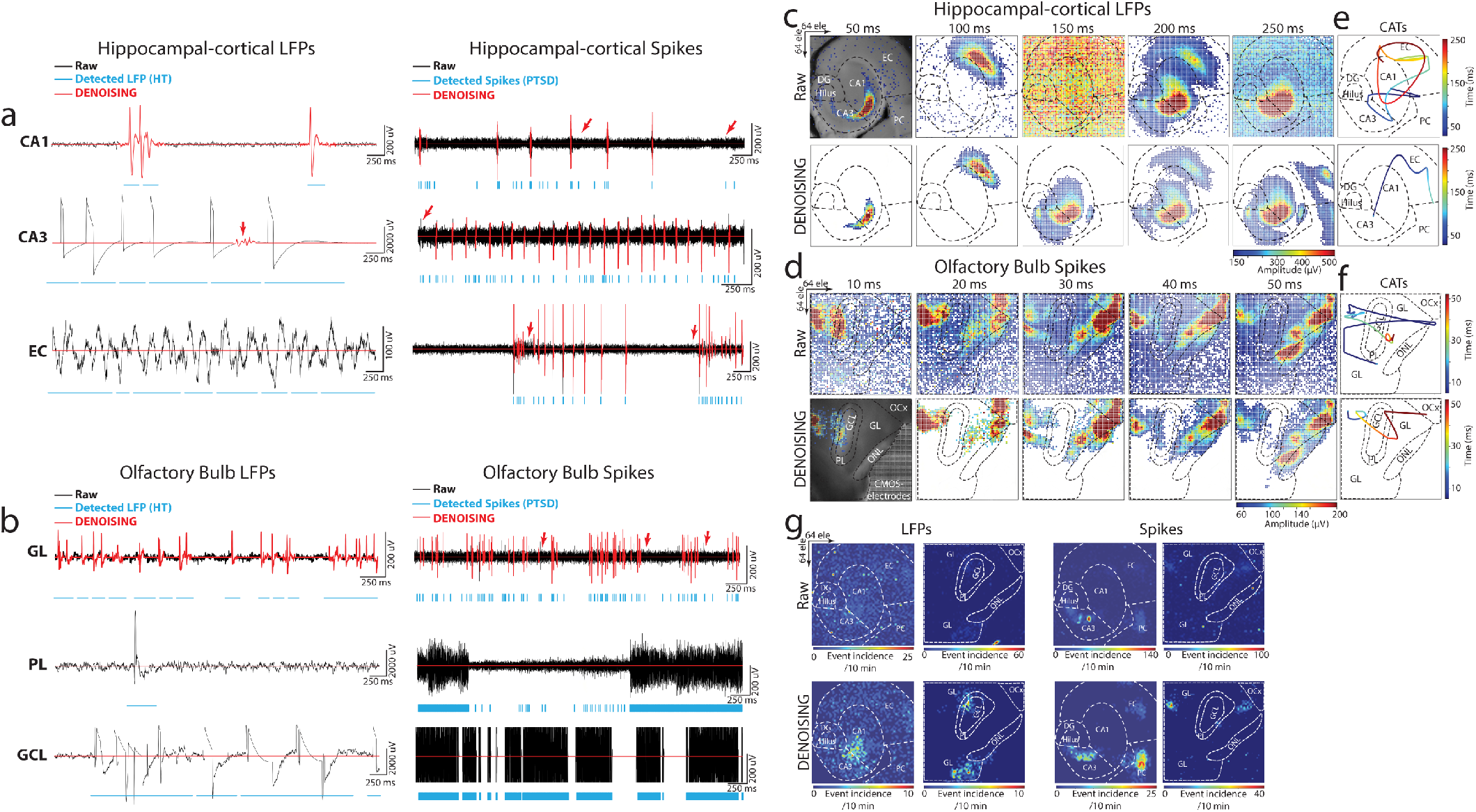
DENOISING Impact on Multimodal Bioelectrical Signal Clarity and Circuit Dynamics. **a-b)** Showcases DENOISING’s precision in hippocampal-cortical and OB circuits, effectively isolating genuine neural activities from noise across CA1, CA3, EC, GL, PL, and GCL regions. Illustrated are examples of successfully discarded spatiotemporal segments of unwanted noise, elimination of non-physiological signals, suppression of inductive coupling noise, and removal of spurious spiking patterns, ensuring fidelity in neural signal interpretation. Red arrows highlight instances where spikes detected by the PTSD algorithm were effectively clarified by DENOISING, showcasing the method’s capability to ensure accurate signal interpretation. **c-d)** Depict the integration of DENOISING-processed neural activity with anatomical landmarks, constructing spatial maps that reveal the organization of neural activity through extracellular discharges in false-color time-lapse frames, highlighting the significant noise reduction achieved compared to raw frames. **e-f)** Center-of-activity trajectories (CATs) from LFP and spiking events delineate information transfer within neural circuits, demonstrating DENOISING’s impact on refining data interpretation by revealing true global circuit dynamics illustrated through the propagation paths of CATs and their timing. **g)** Illustrates the spatial propagation of rhythmic electrical patterns in raw and post-DENOISING, revealing authentic firing patterns and enhancing the fidelity of neural spatiotemporal signal interpretation by elucidating intricate dynamics of neural activity.

Next, to elucidate the profound impact of DENOISING in identifying and mitigating independent noise across the entire spatiotemporal functional landscape within our recorded subnetworks, we integrated large-scale activity with the brain network’s anatomical landmarks. We overlaid the computed mean of firing patterns obtained from LFP and spike recordings onto optical images of hippocampal-cortical and OB circuits; then, we constructed maps that revealed the spatial organization of neural activity (**Figures 2c-d**). These maps encoded extracellular discharges in false-color time-lapse frames depicting full LFP and spiking events. The frames, with time steps of 50 ms for LFP and 10 ms for spiking events, showcased the global noise-embedded contamination across layers in raw data frames compared to the denoised frames. This visualization underscored the substantial reduction in noise achieved by DENOISING, enhancing the clarity and precision of the recorded neural activity. We further constructed center-of-activity trajectories (CATs) from LFP and spiking events to delineate information transfer and processing pathways within the neural circuits. A striking disparity between raw and denoised frames (in events time and amplitude) was observed in the averaged-CAT patterns, emphasizing the pivotal role of DENOISING in refining data interpretation and uncovering the true global circuit dynamics emerging from population activity (**Figures 2e-f**).

Simultaneous recordings from all multimodal network layers were leveraged to compute the generation site and trace the spatial propagation of rhythmic electrical patterns. This analysis illustrated the remarkable significance of DENOISING in providing clarity to unveil the authentic firing patterns obscured by noise in the raw data. The spatial maps of rhythmic electrical patterns elucidated the intricate dynamics of neural activity and highlighted the transformative effect of DENOISING in enhancing the fidelity of neural signal interpretation (**Figure 2g**).

In summary, the integration of robust DENOISING techniques with detailed spatial analyses significantly advances the interpretation of neural activity within key brain networks. This approach aligns functional insights with underlying neural structures, enhancing signal clarity and ensuring that observed behaviors match known anatomical features. This synergy offers crucial insights into complex neural interactions and their physiological implications. These enhanced denoising capabilities could impact the understanding of prolonged depolarization states and the synchronization within neural networks, potentially offering new avenues for research into neural connectivity and its role in cognitive and sensory processing.

### 3.3 DENOISING for Waveform Characterization

Analysis of oscillatory activity waveforms is pivotal for understanding the diverse functionalities within neural circuits. Waveform shapes can indicate different types of neuronal activity and are vital in identifying distinct neuronal types in specific brain regions and their roles in processing information (34). Accurately characterizing waveform features from large-scale neural recordings is essential for decoding the intricate activities within brain networks. This necessitates enhancing signal clarity through meticulous noise reduction, ensuring that the subtle nuances of neural interactions are not lost. To further evaluate the impact of our DENOISING method, we applied PCA and the Mean-Shift algorithm to LFPs extracted from raw and denoised hippocampal and OB recordings. On raw data, PCA depicted a jumbled cluster of signals, indicating a lack of distinct grouping among waveform features. However, post-DENOISING, PCA delineated between groups of data, with color-coded waveforms distinctly belonging to their respective clusters (**Figures 3a-b**). This stark differentiation underscores DENOISING’s capability to significantly enhance the discriminability and clarity of neural signals, allowing for a more nuanced interpretation of complex network activities.

**Figure 3.**
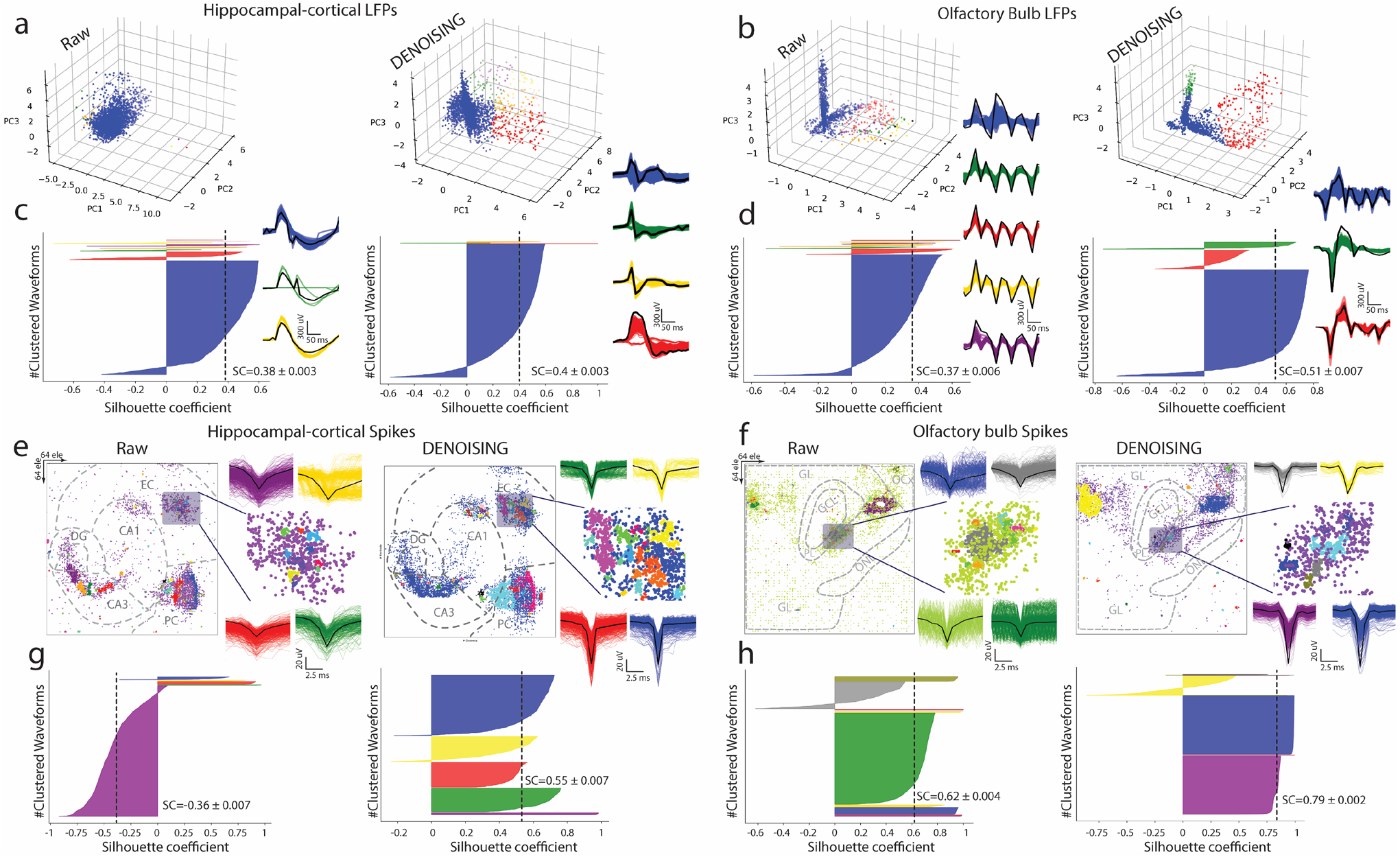
Advanced Waveform Analysis and Clustering Enhancement via DENOISING. **a-b)** Utilizing PCA and Mean-shift algorithm on LFPs from hippocampal-cortical and OB recordings illustrates the transformation from indistinguishable clusters in raw data to clearly defined color-coded waveform groups post-DENOISING, highlighting the method’s success in improving signal discriminability. **c-d)** Silhouette coefficient (SC) analysis further quantifies this clustering enhancement (*p < 0*.*0001 ANOVA*). **e-f)** Advanced spike sorting techniques reveal more distinct spike waveform clusters from hippocampal-cortical and OB recordings. **g-h)** Silhouette coefficients (SC) underscoring the statistical improvement in clustering quality due to DENOISING, emphasizing its pivotal role in refining complex neural signal analysis (*p < 0*.*0001 ANOVA*).

Furthermore, silhouette coefficient analysis quantitatively confirmed the superior clustering performance in denoised data over raw recordings (**Figures 3c-d**).

Expanding our analysis, we employed an advanced spike sorting technique to scrutinize spiking activities within these recordings (30). This process also benefited markedly from the DENOISING process, revealing more defined and distinguishable clusters of spike waveforms. These were coherently matched with their groups identified through principle component (PC) maps, reinforcing the denoising method’s effectiveness (**Figures 3e-f**). The application of silhouette coefficients re-emphasized a statistical foundation for asserting the significant enhancement in clustering quality attributable to our denoising technique, underscoring its effectiveness in refining the analysis of complex neural signals. (**Figures 3g-h**).

### 3.4 Benchmarking with Traditional Noise Removal Approaches

Employing SNR distribution analysis, calculated as the mean over standard deviation, we observed a pronounced enhancement in both LFP and spiking activity data post-DENOISING (**Figures 4a-d**). Specifically, the SNR in hippocampal-cortical and OB LFPs denoised data yielded a 38.8 and 53-fold increase, respectively, over raw data (**Figures 4a-b**). Similarly, spiking activity analysis revealed significant gains, with a 78.8-fold increase in hippocampal-cortical regions and a 16-fold improvement in OB regions (**Figures 4c-d**).

**Figure 4.**
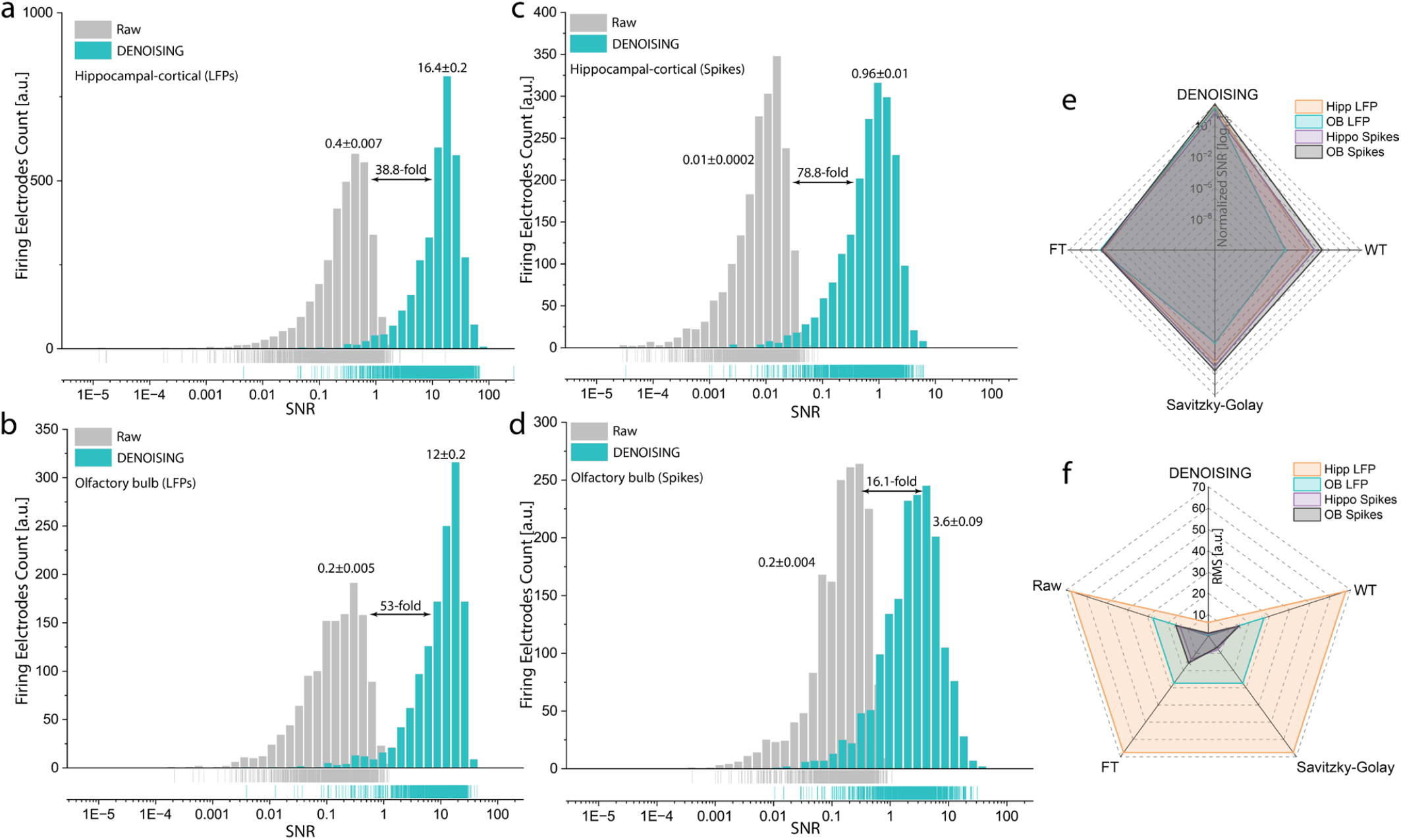
SNR and RMS Enhancements in Neural Signals Post-DENOISING. **a-d)** Depict the significant improvements in SNR distribution for both LFP and spiking activities in hippocampal-cortical and OB recordings following DENOISING application, with significant increases up to 38.8 and 53 times for LFPs, and 78.8 and 16 times for spiking activities, respectively. (LFPs, *p < 10*^*-8*^, *Spikes p < 10*^*-20*^ *Kolmogorov-Smirnov test*). Rug lines under distributions clarify data points by marking their positions on distribution axes, not representing density. **e-f)** Comparative analysis with classical noise reduction techniques through radar plots demonstrates DENOISING’s superior performance, showcasing markedly higher normalized SNR (e) and lower RMS (f) values, thereby affirming its exceptional capability in enhancing signal clarity and accuracy over traditional methods and raw data prior-DENOISING. (SNR LFPs, *p < 10*^*-3*^, SNR Spikes, *p < 10*^*-6*^ *ANOVA*, RMS LFPs, *p < 10*^*-7*^, and RMS Spikes, *p < 10*^*-6*^ *ANOVA*, for DENOISING compared to FT, WT, and Savitzky-Golay).

While classical noise reduction methods have not been explicitly documented in the context of large-scale extracellular recordings and HD-MEAs, we aimed to rigorously evaluate the performance of DENOISING compared to these established techniques – specifically, Wavelet Transform, Fourier Transform, and Savitzky-Golay filter. Our objective was to comprehensively assess DENOISING’s efficacy in enhancing signal clarity and accuracy, which is critical for deciphering the nuanced dynamics of neural activity within the hippocampal-cortical and OB networks.

Further comparative analysis employed the RMS parameter and normalized SNR of denoised data to raw, illustrating DENOISING’s superiority. Radar plots displayed that denoised data exhibited significantly higher SNR (**Figures 4e**) and lower RMS values (**Figures 4f**) for both LFPs and spikes, outperforming classical noise-removal methods. These findings underscore the advanced capability of DENOISING in refining neural signal analysis, setting a new benchmark for the analysis of complex extracellular recordings.

## 4 Discussion

We have introduced DENOISING, a computational framework validated with experimental multimodal data to enhance fidelity and clarity through dynamic noise mitigation on large-scale bioelectrical signals. The adaptive nature of the method allowed for handling a wide range of noise, including electrical, mechanical, and environmental sources, ensuring robust noise removal while preserving the essential features of a diverse array of extracellular LFP and spike signals. This capability is particularly relevant in the context of recent advancements in large-scale biosensing HD-MEAs and their application in monitoring neural dynamics across various spatial and temporal scales. By facilitating precise observations of network-wide activity, our approach enabled a deeper understanding of the interconnected processes governing brain function, which has profound implications for areas critical to learning, memory, and sensory processing.

Central to our results is the improved accuracy in waveform clustering, spike sorting, and CAT analysis, allowing for more precise identification and categorization of neural signals and network dynamics. This precision is critical for understanding the nuanced activities within neural circuits and contributes to a more explicit interpretation of neural communication patterns. Through significant SNR improvements and RMS noise reduction analyses, our methodology provided compelling evidence of its performance that transcends traditional methods. This enhancement not only improved the quality and reliability of neural data but also enabled the detection of subtle neural activity, thus unveiling obscured neural dynamics and interactions that were previously indiscernible.

Furthermore, our study aligns with the growing body of research emphasizing the critical need for high-fidelity neural data to understand complex brain functions and disorders (10,11,35,36), necessitating multi-purpose denoising methods (32,37,38).

In addressing the specific process of our DENOISING method, it is important to clarify that while the algorithm proficiently identifies and improves the clarity of signals associated with significant LFP and spike events, it does not alter or reconstruct signal components outside of these detected events. This is evident in Figure 2, where only meaningful neural activities are highlighted, emphasizing the adaptive nature of our filter in focusing on significant neural events. This selective filtering is vital for accurately interpreting SNR improvements shown in Figure 4, as it directly relates to the assessed signal quality rather than indiscriminately altering the entire data spectrum.

While the DENOISING framework has shown substantial efficacy in enhancing the clarity of extracellular recordings through dynamic adjustment and noise separation, it does not incorporate the application of deep learning approaches exemplified by other methods (32,37,38). However, by not utilizing a machine learning backbone or extensive training datasets, DENOISING offers a significant computational advantage compared to other methods. It requires considerably less computational resources and time, bypassing the need to train a neural network, which is computationally intensive and requires large datasets. This makes DENOISING more accessible for real-time applications and suitable for environments with limited computational capabilities, providing a practical solution for immediate noise reduction without the overhead of training and model optimization. However, integrating a deep learning component into our framework represents a compelling avenue for future research, which could potentially offer improved adaptability and precision in noise reduction and signal processing by harnessing the power of large-scale neural datasets for training purposes.

## 5 Data Availability Statement

The data supporting our results are comprehensively presented within the main text and figures of this article. Data samples with Python script for the method are available on our Lab’s GitHub (https://github.com/HayderAminLab/DENOISING).

## 6 Ethics Statement

All experiments in this study were conducted in accordance with the applicable European and national regulations (Tierschutzgesetz) and were approved by the local authority (Landesdirektion Sachsen; 25– 5131/476/14).

## 7 Author Contributions

X.H. wrote the code and analyzed the multiscale multimodal data. B.A.E. and S.K. performed multimodal extracellular recording experiments. H.A. conceptualized, planned, and supervised the study, designed and performed experiments, developed computational tools, and generated the figures. H.A. wrote the manuscript. All authors reviewed and approved the final manuscript.

## 9 Acknowledgments

We want to acknowledge the support of the platform for behavioral animal testing at the DZNE-Dresden (Dr. Alexander Garthe, Anne Karasinsky, Sandra Günther, and Jens Bergmann).

## REFERENCES

1. Vyas S, Golub MD, Sussillo D, Shenoy K V. Computation through Neural Population Dynamics. Annu Rev Neurosci. 2020;43:249–75.

2. Bird CM, Burgess N. The hippocampus and memory: insights from spatial processing. Nat Rev Neurosci [Internet]. 2008 Mar [cited 2022 Jan 13];9(3):182–94. Available from: https://pubmed.ncbi.nlm.nih.gov/18270514/

3. Mori K, Nagao H, Yoshihara Y. The olfactory bulb: Coding and processing of odor molecule information. Science (1979). 1999;286(5440):711–5.

4. Lisman J, Buzsáki G, Eichenbaum H, Nadel L, Rangananth C, Redish AD. Viewpoints: how the hippocampus contributes to memory, navigation and cognition. Nat Neurosci [Internet]. 2017 [cited 2022 Jan 13];20(11):1434–47. Available from: https://pubmed.ncbi.nlm.nih.gov/29073641/

5. Kempermann G, Gage FH, Aigner L, Song H, Curtis MA, Thuret S, et al. Human Adult Neurogenesis: Evidence and Remaining Questions. Cell Stem Cell. 2018;23(1):25–30.

6. Lepousez G, Valley MT, Lledo PM. The Impact of Adult Neurogenesis on Olfactory Bulb Circuits and Computations. Annu Rev Physiol. 2013 Feb 10;75(1):339–63.

7. Buzsáki G, Anastassiou CA, Koch C. The origin of extracellular fields and currents-EEG, ECoG, LFP and spikes. Nat Rev Neurosci. 2012;13(6):407–20.

8. Berdondini L, Imfeld K, Maccione A, Tedesco M, Neukom S, Koudelka-Hep M, et al. Active pixel sensor array for high spatio-temporal resolution electrophysiological recordings from single cell to large scale neuronal networks. Lab Chip [Internet]. 2009 Sep 21 [cited 2015 Feb 3];9(18):2644–51. Available from: http://pubs.rsc.org/en/content/articlehtml/2009/lc/b907394a

9. Müller J, Ballini M, Livi P, Chen Y, Radivojevic M, Shadmani A, et al. High-resolution CMOS MEA platform to study neurons at subcellular, cellular, and network levels. Lab Chip. 2015;15(13):2767–80.

10. Hu X, Khanzada S, Klütsch D, Calegari F, Amin H. Implementation of biohybrid olfactory bulb on a high-density CMOS-chip to reveal large-scale spatiotemporal circuit information. Biosens Bioelectron [Internet]. 2022;198(March 2021):113834. Available from: 10.1016/j.bios.2021.113834

11. Emery BA, Hu X, Khanzada S, Kempermann G, Amin H. High-resolution CMOS-based biosensor for assessing hippocampal circuit dynamics in experience-dependent plasticity. Biosens Bioelectron [Internet]. 2023;237(May):115471. Available from: 10.1016/j.bios.2023.115471

12. Amin H, Maccione A, Marinaro F, Zordan S, Nieus T, Berdondini L. Electrical responses and spontaneous activity of human iPS-derived neuronal networks characterized for 3-month culture with 4096-electrode arrays. Front Neurosci. 2016;10(121):1–15.

13. Amin H, Nieus T, Lonardoni D, Maccione A, Berdondini L. High-resolution bioelectrical imaging of Aβ-induced network dysfunction on CMOS-MEAs for neurotoxicity and rescue studies. Sci Rep [Internet]. 2017 Dec 26 [cited 2017 May 29];7(1):2460. Available from: http://www.ncbi.nlm.nih.gov/pubmed/28550283

14. Amin H, Marinaro F, Tonelli DDP, Berdondini L. Developmental excitatory-to-inhibitory GABA-polarity switch is disrupted in 22q11.2 deletion syndrome: A potential target for clinical therapeutics. Sci Rep [Internet]. 2017;7(1):1–18. Available from: 10.1038/s41598-017-15793-9

15. Buzsáki G. Large-scale recording of neuronal ensembles. Nat Neurosci [Internet]. 2004 May;7(5):446–51. Available from: 10.1038/nn1233

16. Rossi L, Emery BA, Khanzada S, Hu X, Amin H. Pharmacologically and Electrically-induced Network-wide Activation of Olfactory Bulb with Large-scale Biosensor. In: 2023 IEEE BioSensors Conference (BioSensors) [Internet]. IEEE; 2023. p. 1–4. Available from: https://ieeexplore.ieee.org/document/10280876/

17. Emery BA, Hu X, Maugeri L, Khanzada S, Klütsch D, Amin H. Large-scale Multimodal Neural Recordings on a High-density Neurochip: Olfactory Bulb and Hippocampal Networks. IEEE EMBS. 2022;42–5.

18. Emery BA, Khanzada S, Hu X, Rossi L, Klütsch D, Altuntac E, et al. Recording Network-based Synaptic Transmission and LTP in the Hippocampal Network on a Large-scale Biosensor. In: 2023 IEEE BioSensors Conference (BioSensors) [Internet]. IEEE; 2023. p. 1–4. Available from: https://ieeexplore.ieee.org/document/10280958/

19. Harris KD, Henze DA, Csicsvari J, Hirase H, Buzsáki G. Accuracy of tetrode spike separation as determined by simultaneous intracellular and extracellular measurements. J Neurophysiol. 2000;84(1):401–14.

20. Patil R. Noise Reduction using Wavelet Transform and Singular Vector Decomposition. Procedia Comput Sci [Internet]. 2015;54:849–53. Available from: 10.1016/j.procs.2015.06.099

21. Donoho DL. De-Noising by Soft-Thresholding. 1995;41(3).

22. Starck JL, Candès EJ, Donoho DL. The curvelet transform for image denoising. IEEE Transactions on Image Processing. 2002;11(6):670–84.

23. Kay K. The risk of bias in denoising methods: Examples from neuroimaging. PLoS One [Internet]. 2022;17(7 July):1–19. Available from: 10.1371/journal.pone.0270895

24. Czanner G, Sarma S V., Ba D, Eden UT, Wu W, Eskandar E, et al. Measuring the signal-to-noise ratio of a neuron. Proc Natl Acad Sci U S A. 2015;112(23):7141–6.

25. Gustafsson F. Determining the initial states in forward-backward filtering. IEEE Transactions on Signal Processing. 1996;44(4):988–92.

26. Krishnan SR, Seelamantula CS. On the selection of optimum Savitzky-Golay filters. IEEE Transactions on Signal Processing. 2013;61(2):380–91.

27. Minka TP. Automatic dimensionality selection for PCA. 2008;

28. Comaniciu D, Meer P. Mean shift: A robust approach toward feature space analysis. IEEE Trans Pattern Anal Mach Intell. 2002;24(5):603–19.

29. Coates A, Ng AY. Learning Feature Representations with K-means. Lecture Notes in Computer Science (including subseries Lecture Notes in Artificial Intelligence and Lecture Notes in Bioinformatics). 2012;7700 LECTU:561–80.

30. Hilgen G, Sorbaro M, Pirmoradian S, Muthmann JO, Kepiro IE, Ullo S, et al. Unsupervised Spike Sorting for Large-Scale, High-Density Multielectrode Arrays. Cell Rep [Internet]. 2017;18(10):2521–32. Available from: 10.1016/j.celrep.2017.02.038

31. Rousseeuw PJ. Silhouettes: A graphical aid to the interpretation and validation of cluster analysis. J Comput Appl Math. 1987;20(C):53–65.

32. Lecoq J, Oliver M, Siegle JH, Orlova N, Ledochowitsch P, Koch C. Removing independent noise in systems neuroscience data using DeepInterpolation. Nat Methods. 2021;18(11):1401–8.

33. Hyndman RJ, Koehler AB. Another look at measures of forecast accuracy. Int J Forecast. 2006;22(4):679–88.

34. Cole SR, Voytek B. Brain Oscillations and the Importance of Waveform Shape. Trends Cogn Sci. 2017;21(2):137–49.

35. Li H, Wang J, Fang Y. Recent developments in multifunctional neural probes for simultaneous neural recording and modulation. Microsyst Nanoeng. 2023;9(1).

36. Vázquez-Guardado A, Yang Y, Bandodkar AJ, Rogers JA. Recent advances in neurotechnologies with broad potential for neuroscience research. Nat Neurosci. 2020;23(12):1522–36.

37. Eom M, Han S, Park P, Kim G, Cho ES, Sim J, et al. Statistically unbiased prediction enables accurate denoising of voltage imaging data. Nat Methods. 2023;20(10):1581–92.

38. Li X, Zhang G, Wu J, Zhang Y, Zhao Z, Lin X, et al. Reinforcing neuron extraction and spike inference in calcium imaging using deep self-supervised denoising. Nat Methods. 2021;18(11):1395–400.

